# Early resolution of sister chromatids during *C. elegans* meiosis

**DOI:** 10.1101/2025.03.29.646130

**Authors:** Antonia Hamrick, Ofer Rog

## Abstract

Segregating a complete set of chromosomes into the gametes relies on exchanges of genetic material that occur during meiosis. It is only exchanges that form between the similar parental chromosomes (homologs), rather than between the identical sister chromatids, that enable correct chromosome segregation (Zickler and Kleckner 2023). Despite the crucial role of biasing exchanges toward the homolog and the progress in defining some of its regulators (Niu *et a*l. 2009; Goldfarb and Lichten 2010; Lao and Hunter 2010; Kim *et al*. 2010; Hong *et al*. 2013), the mechanism that efficiently identifies the homolog and avoids the sister remains unknown. Understanding homolog bias requires knowledge of how the homologs and sisters are organized relative to each other, and how this positioning is established. Here, we use selective labeling of a single sister in the oogenic germline of the nematode *Caenorhabditis elegans* to define the organization of the sister chromatids at the time exchanges form. We find that pairs of sisters are already well separated (resolved) early in meiosis, despite being tethered to each other at numerous positions along their length. The sisters are resolved in both aligned and unaligned homologs, and their resolution does not require condensins or a prolonged time in meiotic prophase. However, depleting the cohesin loader NIPBL^SCC-2^ impairs sister resolution, suggesting that an active process - likely loop extrusion by cohesins - de-mixes and resolves the sisters. Our work shows that inter-homolog meiotic exchanges form when the four sister chromatids occupy distinct volumes, suggesting that homolog bias is unlikely to rely on relative proximity. The conservation of meiotic chromosome organization and of cohesin’s loop-extruding activity suggests that our findings are broadly applicable.

## Results and Discussion

### Sister chromatids are resolved during meiotic prophase

Nuclei in the *C. elegans* gonad progress through the different stages of meiosis as they move through the gonad. Upon their exit from a stem-cell-like compartment, meiotic nuclei replicate their DNA, organize their chromatin as an array of loops tethered at their base to a proteinaceous structure that organizes them (called the ‘axis’), and align their homologs. The aligned homologs undergo double-stranded breaks (DSBs), a subset of which is repaired to form the inter-homolog exchanges (crossovers) that subsequently allow them to segregate into the gametes (Fig. S1; (Zickler and Kleckner 2023)). Whereas the side-by-side organization of the homologs has been characterized in diverse organisms, the organization of the two sisters within each homolog is poorly understood due to their identical sequence. We adapted our recently developed EdU labeling protocol (STAR Methods; (Almanzar *et al*. 2021, 2022)) so that one of each pair of sister chromatids is labeled at the time when DSBs are being repaired. Together with the superior cytological properties of the worm germline (Phillips *et al*. 2009), this approach allowed us to define the organization of identical pairs of sister chromatids in structurally preserved meiotic nuclei.

We used 3D-STED microscopy to analyze sister organization in wild-type worms. We readily observed incomplete overlap between the EdU signal, which marks one sister, and the DNA signal, which marks both sisters (‘single-labeled’; Fig. 1A). We also observed incomplete overlap when imaging our samples with 2D-STED microscopy, as well as when using a different dye combination to label EdU and DNA, arguing against optical artifacts (Fig. S2A). Importantly, we observed an almost complete overlap between the EdU and DNA signals when we adjusted the EdU incorporation protocol so that both sisters were labeled (referred to throughout as ‘double-labeled’; Fig. 1A and S2A; STAR Methods).

**Figure 1:**
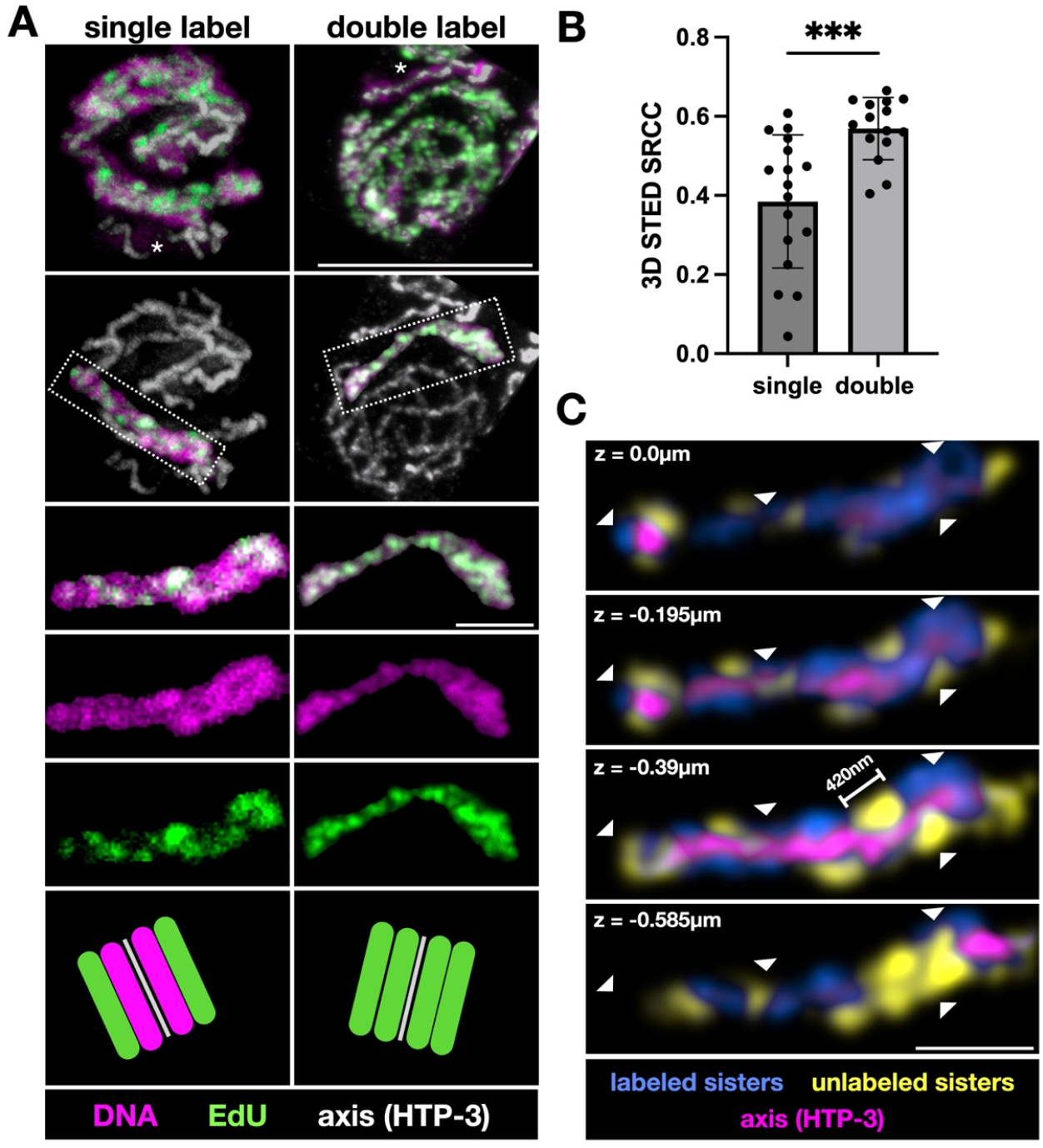
Sister chromatids are resolved during meiotic prophase. A. 3D-STED images of nuclei with single- and double-labeled chromosomes. Top row, projections of whole nuclei. DNA (LIVE460L), purple; EdU (Alexa594), green; axis (HTP-3), white. Asterisks represent an X chromosome that is unlabeled due to its late replication (Jaramillo-Lambert *et al*. 2007). Second row, a single chromosome; for clarity, the DNA and EdU signals were computationally removed from the other chromosomes. Third row, merged images of a single chromosome. Fourth and fifth rows, single channel images of a single chromosome. Scale bars are 5μm for the whole nucleus and 1μm for the single chromosome. Bottom, diagram of single- and double-labeled chromosomes, with each colored bar representing a single sister chromatid. B. Spearman’s rank correlation coefficient (SRCC) between the DNA and EdU channels from single- and double-labeled chromosomes (3D-STED images; n=17 and 15, respectively). ANOVA p=0.0001. C. Single z-sections of the indicated depth from 3D-STED microscopy, with the labeled and unlabeled sisters in blue and yellow, respectively, and the axis (HTP-3) in magenta. The unlabeled sisters (yellow) were computationally derived. Arrowheads indicate labeled and unlabeled clusters of chromatin. Scale bar = 1μm.

To quantify the degree of overlap, we calculated Spearman’s rank correlation coefficient (Batty *et al*. 2023). In this statistical modality, all pixels in a region of interest (in our case, an aligned chromosome pair) are ranked according to their intensity, and then their rank in each channel is compared. Perfectly overlapping signals yield a correlation coefficient of 1, while resolved sisters are expected to yield a lower value. An idealized scenario of completely resolved sisters, where half of the DNA pixels harbor an EdU signal of similar intensity, will yield a correlation coefficient of 0.5. When both sisters were labeled, we found a correlation coefficient of 0.56. Minor optical aberrations and low signal likely account for this number being lower than the expected correlation coefficient of 1. Importantly, however, we found a significantly lower correlation for single-*versus* double-labeled sisters, 0.38 *versus* 0.56 (unpaired t-test, p<0.005; Fig. 1B). Normalizing these values under the assumption of an underlying complete overlap in the double-label scenario (Batty *et al*. 2023), yields a correlation coefficient of 0.67 for the single-label sister, not far from the expected value of 0.5 for complete sister resolution. These data indicate that the incomplete overlap between the DNA and EdU signals reflects *bone fide* resolution of the sisters during pachytene, the meiotic substage when crossovers form.

To allow a better understanding of the morphology of the sisters, we computationally derived a visualization of the unlabeled sister (Fig. 1C; STAR Methods). Serial 3D-STED z-sections revealed 300-500nm clusters of chromatin that are organized on top of or next to each other, suggesting that each sister chromatid forms a thick cylinder, with each pair sisters braided along their length. This morphology is consistent with fluorescence *in situ* hybridization (FISH) analysis of a single locus, which revealed that chromatin loops belonging to pairs of sisters are commonly found separated (Woglar *et al*. 2020). The 300-500nm EdU-positive and EdU-negative domains (Fig. 1C) contain ∼1Mbp of DNA derived from one of the co-linear sisters (based on a ∼2.5Mbp/µm packaging of meiotic chromosomes; (Albertson *et al*. 1997)). The estimated average size of chromatin loops - ∼100-150kbps (Woglar *et al*. 2020) - suggests that adjacent chromatin loops belonging to the same sister are clustered and in close proximity.

To address the potential role of homolog alignment in sister resolution, we analyzed a condition under which only two sisters are tethered to one another. We used worms lacking *him-8*, where the X chromosomes fail to pair and align, although they undergo DSB formation and repair (Phillips *et al*. 2005). Single sister labeling and imaging by confocal microscopy revealed incomplete overlap between the EdU and DNA signals (Fig. 2A), similar to our observations in paired and aligned homologs above.

**Figure 2:**
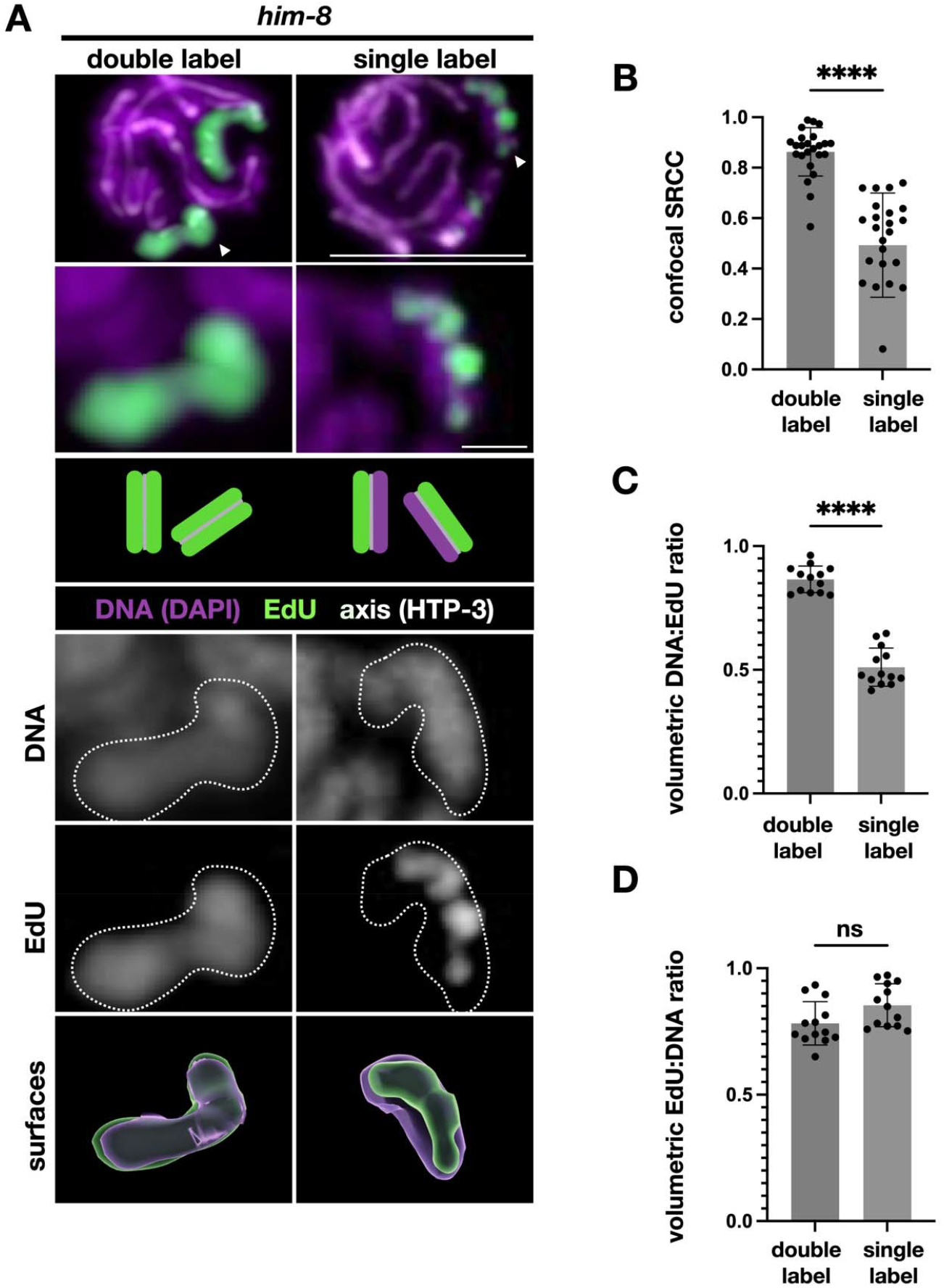
Sister chromatids in unaligned homologs are resolved during meiotic prophase. A. Confocal images of nuclei from *him-8* animals with single- and double-labeled chromosomes where only the X chromosomes (white arrowheads) are labeled. DNA (DAPI), magenta; EdU, green; axis (HTP-3), white. Second row, zoomed-in view of the unaligned EdU-labeled X chromosome, with diagrams of the labeling scheme below. Third and fourth row, single-channel greyscale images, with the outline of the chromosome shown (dashed white line). Bottom, constructed volumetric surfaces shown in transparent green and magenta for EdU and DAPI, respectively. Scale bar is 5μm for the whole nucleus and 1μm for the single chromosome. B. Spearman’s rank correlation coefficient (SRCC) between the DNA and EdU channels from confocal images of double- and single-label unaligned chromosomes in *him-8* worms (n=24 for each sample). Unpaired t-test p<0.0001. C. Ratio of DNA to EdU of constructed surfaces for *him-8* X chromosome double- and single-labeled chromosomes (n=13 for each sample). ANOVA p<0.0001. D. Ratio of EdU to DNA, as in panel B. ANOVA p=0.10.

Spearman’s rank correlation coefficient between the EdU and DNA channels was 0.49 and 0.86 for single- and double-labeled unaligned X chromosomes, respectively (Fig. 2D; unpaired t-test, p<0.0001). We also observed a low correlation between the DNA and EdU channels in 2D-STED data (Fig. S2A-B). The low correlation in the single-labeled nuclei suggests that the sisters are well resolved, even at the resolution examined here (for confocal microscopy, approximately 150×150×500nm). The more easily apparent spatial separation of the sisters in unaligned (Fig. 2) *versus* aligned (Fig. 1) homologs suggests that the presence of two aligned homologs may limit sister resolution, potentially due to the space occupied by the homolog and the more limited chromatin mobility around the axes. However, other differences in morphology and chromatin state between aligned and unaligned homologs may underlie this difference (Lui and Colaiácovo 2013; Yu *et al*. 2016; Zickler and Kleckner 2023).

The relative ease of interpreting sister organization in unaligned homologs allowed us to use volumetric analysis as an orthogonal way to quantify sister resolution. We used 3-dimensional confocal data to construct two volumetric surfaces based on the DNA and EdU signals of the X chromosome in *him-8* worms (Fig. 2A, bottom; STAR Methods). We found that 51% of the DNA surface is overlapping with the EdU surface (Fig. 2B). This DNA:EdU ratio reflects the theoretical maximum separation between two identical objects, indicating that the sisters mostly occupy non-overlapping volumes and are essentially fully resolved. As expected, the inverse ratio between the same surfaces - the amount of EdU surface that overlaps with the DNA surface - was 85%, close to the expected 100%. As a control, we quantified sister resolution when both sisters within each chromosome were labeled and found a significantly higher overlap - DNA:EdU ratio of 87% (unpaired t-test, p<0.0001; Fig. 2C) - further validating our volumetric quantification.

### Sisters resolve early during meiosis

The meiotic cell cycle is much longer than its mitotic counterpart, and lasts, in worms, about 24 hours (nuclei progress in the gonad at a rate of ∼1 nuclei-row / hour; Fig. S1; (Crittenden *et al*. 2006; Fox *et al*. 2011)). We wondered whether sister resolution is a consequence of the extended duration - 15-20 hours - between premeiotic S-phase and pachytene, the meiotic stage that was analyzed above (Figs. 1-2). We therefore examined nuclei about 5 hours after premeiotic S-phase, in early pachytene - the time when meiotic DSBs are first introduced (∼10 hours earlier in meiosis compared with our analysis above; Fig. 3A; (Colaiácovo *et al*. 2003)). Despite overall nuclear morphology being less organized in early pachytene (Carlton *et al*. 2006), we found a mean DNA:EdU ratio of 48% - not significantly different from the nuclei analyzed in Fig. 2 (unpaired t-test, p=0.42; Fig. 3B). This indicates that the sisters are resolved throughout the window of meiotic DSB formation and repair. Furthermore, this observation suggests that sister resolution is not merely a consequence of the very long duration of meiotic prophase.

**Figure 3:**
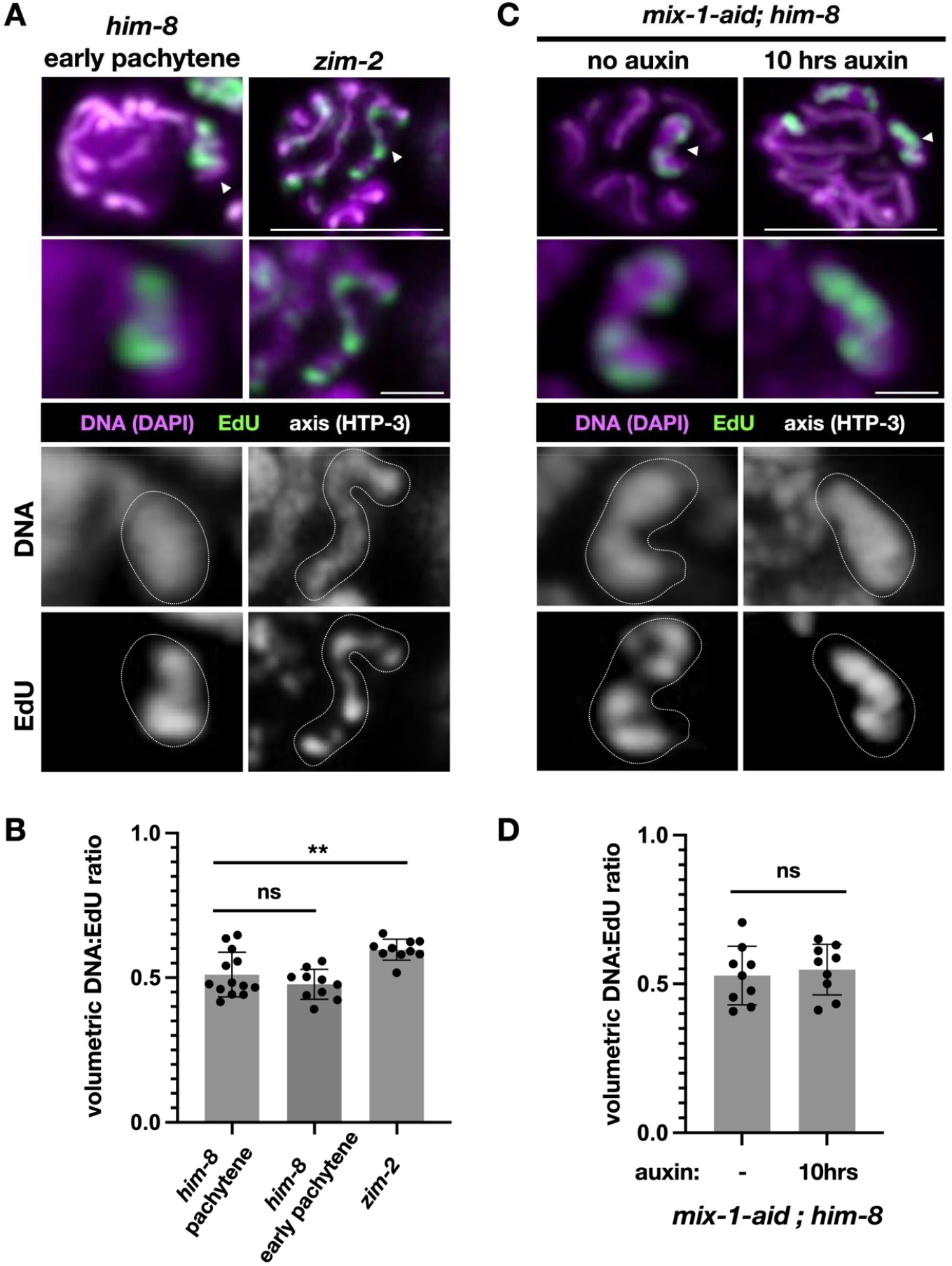
Characterization of sister resolution early in meiosis, on autosomes and sex chromosomes and upon condensin depletion. A. Top, confocal images of pachytene nuclei from *him-8* and *zim-2* worms. DNA (DAPI), purple; EdU, green; axis (HTP-3), white. Second row, zoomed-in view of the unaligned, EdU-labeled chromosome. The X chromosome is unaligned in *him-8* animals and chromosome V is unaligned in *zim-2* animals. Third and fourth rows, single-channel greyscale images, with the outline of the chromosome shown (dashed white line). Note the elongated morphology of chromosome V relative to the X chromosome (Fig. S2D). Scale bar is 5μm for the whole nucleus and 1μm for the single chromosome. B. Volumetric DNA:EdU ratio of *him-8* early pachytene and *zim-2*, compared with *him-8* pachytene from Fig. 2A (n=10, 10 and 13, respectively). See Fig. S2C for the inverse volumetric EdU:DNA ratios. ANOVA *him-8* pachytene vs *him-8* early pachytene p=0.52. ANOVA *him-8* pachytene vs *zim-2* p=0.0019. C. Top, confocal images of nuclei from *mix-1-aid; him-8* worms without auxin and after 10 hours on auxin. This strain also expresses the ubiquitin E3 ligase TIR1 in the germline, which degrades proteins with an AID degron upon exposure to auxin. DNA (DAPI), purple; EdU, green; axis (HTP-3), white. Second row, zoomed-in view of the unpaired, EdU-labeled X chromosome. Third and fourth rows, single-channel greyscale images, with the outline of the X chromosome shown (dashed white line). Scale bar is 5μm for the whole nucleus and 1μm for the single chromosome. D. Volumetric DNA:EdU ratio of *mix-1-aid; him-8* worms without and with auxin treatment (n=9 for both samples). See Fig. S3C for the inverse volumetric EdU:DNA ratios. Unpaired t-test p=0.65.

### Sister resolution in both X chromosomes and autosomes

The X chromosome is regulated differently from the autosomes during meiosis (Kelly *et al*. 2002; Jaramillo-Lambert *et al*. 2007; Wagner *et al*. 2010; Mlynarczyk-Evans and Villeneuve 2017), potentially due to its distinct chromatin composition, which bears some resemblance to classical heterochromatin (e.g., generally repressed transcription; (Kelly *et al*. 2002; Gerstein *et al*. 2010)). To confirm that sister resolution is not specific to the X chromosome, we analyzed *zim-2* worms, where it is chromosome V that does not pair and align with its homolog (Phillips and Dernburg 2006). We found incomplete overlap between the EdU and DNA signals (Fig. 3A), yielding DNA:EdU ratio of 59% (Fig. 3B), indicating that the chromosome V sisters are also resolved. While chromatin composition may underlie the small difference in the degree of sister resolution between the unaligned X chromosomes and the unaligned chromosome V (51% *versus* 59%, respectively; unpaired t-test, p=0.004), it is important to note that chromosome V adopts a more elongated morphology during meiosis (Fig. S2D), which may have influenced our quantification of sister resolution.

### Sister resolution does not require condensin function

Condensins are crucial for sister hyper-condensation and for their resolution just prior to chromosome segregation in mitosis and meiosis (Yatskevich *et al*. 2019). We wished to test whether condensins are also required for sister resolution in meiotic prophase. Since condensins are essential for viability and germline proliferation, we conditionally depleted Smc2^MIX-1^, a component of all three condensin complexes in worms (Csankovszki *et al*. 2009). We inserted an AID degron on the C-terminus of Smc2^MIX-1^, allowing us to conditionally degrade Smc2^MIX-1^ upon exposure to auxin (Zhang *et al*. 2015). This allele exhibited no meiotic defects in the absence of auxin. However, after 24 hours on auxin, the germline exhibited gross abnormalities consistent with a lack of condensin function (Fig. S3). Ten hours of auxin exposure recapitulated the phenotypes of condensin hypomorphs (Mets and Meyer 2009) but did not disrupt the morphology of pachytene nuclei (Fig. S3). In Smc2^MIX-1^-depleted worms that also lack HIM-8, the sisters in the unaligned X chromosomes were resolved (Fig. 3C), exhibiting a similar DNA:EdU ratio to no auxin controls, 55% and 53%, respectively (unpaired t-test, p=0.08; Fig. 3D).

While we cannot rule out a role for the residual amount of condensin in Smc2^MIX-1^-depleted worms, our data suggest that condensin function is not essential for sister resolution in meiotic prophase. Interestingly, condensin depletion did not alter the degree of overlap between the sisters in G2 human somatic cells (Batty *et al*. 2023). These findings echo ours, given that the nuclei we analyzed are undergoing abundant transcription and DNA repair, resembling G2 cells despite being in meiotic prophase.

### Sister resolution requires loop extrusion by cohesin

Cohesins contribute to chromosome dynamics by carrying out two key molecular functions: 1) tethering two DNA molecules (“cohesion”) and 2) loop extrusion, an ATP-dependent motor activity that pulls on the DNA to create progressively enlarging loops (Yatskevich *et al*. 2019; Davidson *et al*. 2019; Kim *et al*. 2019; Castellano-Pozo *et al*. 2023). Polymer simulations have proposed that the latter function can promote sister resolution (Goloborodko *et al*. 2016). During meiosis in worms, these functions are thought to be carried out by two distinct cohesin complexes, named after their kleisin subunits: REC-8 cohesins carry out cohesion by tethering the sisters in numerous positions along their length, while COH-3/4 cohesins perform loop extrusion (COH-3 and -4 are redundant with each other; (Tzur *et al*. 2012; Severson and Meyer 2014; Woglar *et al*. 2020; Castellano-Pozo *et al*. 2023)). To carry out both functions, cohesins must first be loaded onto chromosomes by NIPBL^SCC-2^, with REC-8 cohesins loaded during premeiotic S-phase and COH-3/4 cohesins continuously loaded throughout meiotic prophase (Lightfoot *et al*. 2011; Castellano-Pozo *et al*. 2023).

To address the potential role of loop extrusion in sister resolution, we conditionally depleted NIPBL^SCC-2^ and analyzed sister resolution in the unaligned X chromosomes (*scc-2-aid; him-8* worms; (Castellano-Pozo *et al*. 2023)). We identified a duration - 10 hours on auxin - that depleted NIPBL^SCC-2^ without disrupting nuclear morphology in pachytene (Fig. S4). NIPBL^SCC-2^ depletion led to significantly more overlap between the EdU and DNA signals in unaligned X chromosomes. We found a DNA:EdU ratio of 81% in auxin treated (NIPBL^SCC-2^-depleted) animals *versus* 59% in no auxin controls (Fig. 4A-B; unpaired t-test, p<0.0001). Analysis of aligned chromosomes using 3D-STED (Fig. 4C) similarly revealed more overlap between the EdU and DNA signals in NIPBL^SCC-2^-depleted animals compared to wild-type animals, with Spearman’s ranked correlation coefficient of 0.60 *versus* 0.38 in wild-type worms (p<0.0001; Fig. 4D). These observations indicate that cohesin loading by NIPBL^SCC-2^ is necessary for sister resolution during meiotic prophase, as was found for sister overlap in somatic human cells (Batty *et al*. 2023).

**Figure 4:**
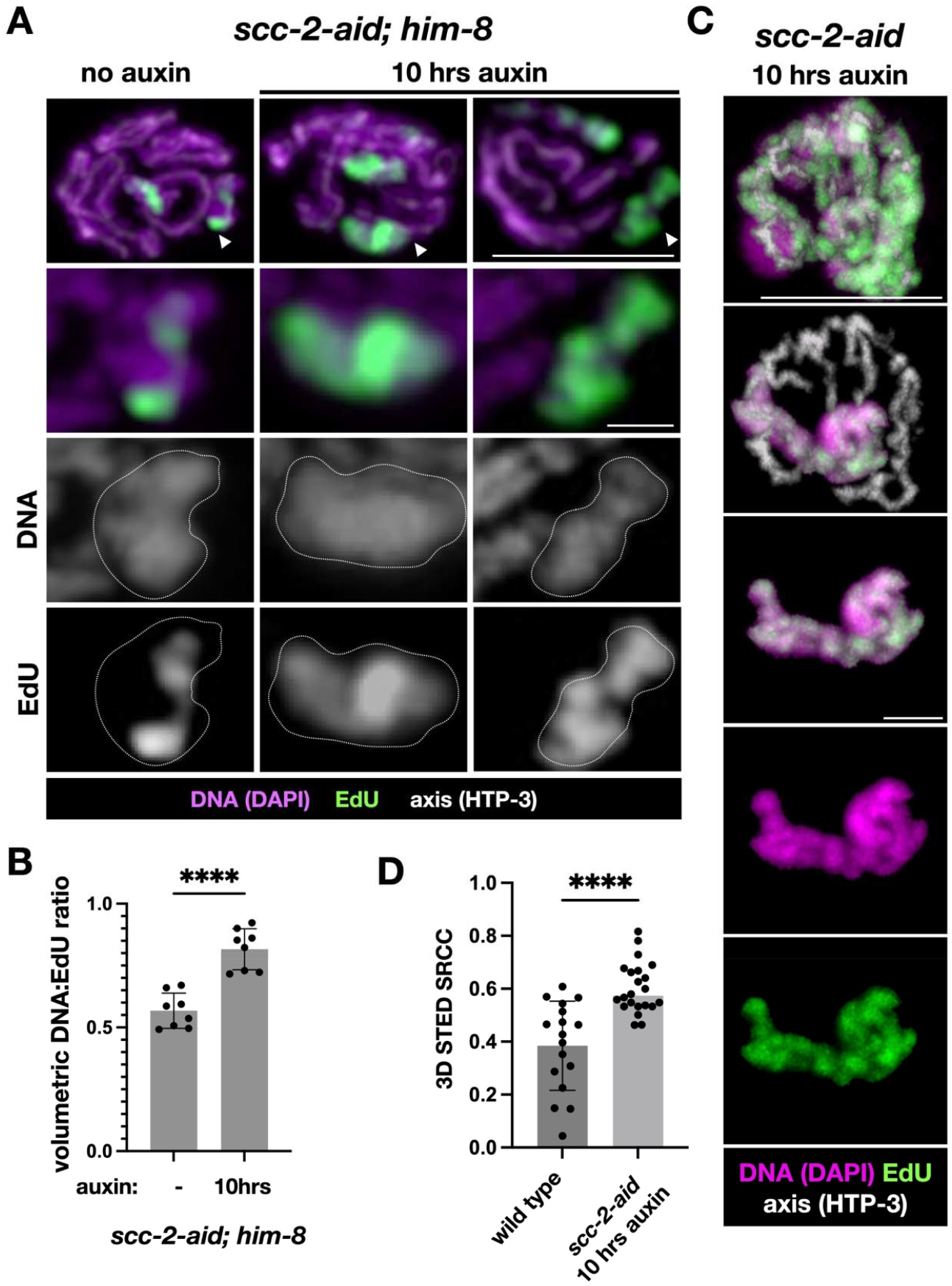
NIPBL^SCC-2^ promotes sister resolution. A. Top, c *scc-2-aid; him-8* worms without auxin or after 10 hours on auxin. This strain also expresses the ubiquitin E3 ligase TIR1 in the germline, which degrades proteins with an AID degron upon exposure to auxin. DNA (DAPI), purple; EdU, green; axis (HTP-3), white. White arrowheads denote the EdU-labeled unpaired X chromosome. Second row, zoomed in view of the unaligned EdU labeled X chromosome. Third and fourth row, single-channel greyscale images, with the outline of the X chromosome shown (dashed white line). Scale bar is 5μm for the whole nucleus and 1μm for the single chromosome. B. Volumetric DNA:EdU ratio of worms without and with auxin (n=8 for each sample). Unpaired t-test p<0.0001. See Fig. S4C for the inverse EdU:DNA ratios. C. Top, 3D-STED image of a single-labeled nucleus from *scc-2-aid* animals treated with auxin for 10 hours. DNA (LIVE460L), purple; EdU (Alexa594), green; axis (HTP-3), white. Second row, a single chromosome; for clarity, the DNA and EdU signals were computationally removed from the other chromosomes. Third row, merged images of a single chromosome. Fourth and fifth rows, single channel images of a single chromosome. Scale bars are 5μm for the whole nucleus and 1μm for the single chromosome. D. Spearman’s rank correlation coefficient (SRCC) between the DNA and EdU channels (3D-STED) of *scc-2-aid* animals treated with auxin for 10 hours compared to wild-type controls (from Fig. 1B; n=22 and 17, respectively). ANOVA p< 0.0001.

The NIPBL^SCC-2^-depleted nuclei analyzed above underwent premeiotic DNA replication - the time when REC-8 cohesin is loaded - prior to NIPBL^SCC-2^ being degraded (Fig. S1 and S4; Castellano-Pozo *et al*. 2023)). This suggested that REC-8 cohesin may not be essential for sister resolution. Consistent with this idea, analysis of *rec-8* worms revealed that sisters are still resolved, with a mean DNA:EdU ratio of 49%, indistinguishable from *him-8* animals (Fig. S5). (Note that in *rec-8* worms all homologs fail to align (Pasierbek *et al*. 2001; Cahoon *et al*. 2019), allowing us to compare them to the unaligned X chromosomes in *him-8* worms; in addition, based on their late replication timing, 10/11 chromosomes analyzed in *rec-8* worms were in fact unaligned X chromosomes.)

We next tested whether COH-3/4 cohesin plays a role in sister resolution given its continuous NIPBL^SCC-2^-dependent loading throughout meiotic prophase and proposed loop extruding activity (Castellano-Pozo *et al*. 2023). We constructed a *coh-3-aid* allele in worms that also lack *coh-4* and identified a duration of auxin exposure that conferred depletion of COH-3 with minimal effect on nuclear morphology in pachytene (Fig. S6). Unexpectedly, analysis of sister resolution in *coh-3-aid coh-4; him-8* worms placed on auxin for 10 hours revealed mean DNA:EdU overlap of 56%, similar to control *him-8* worms (Fig. S5). We hypothesize that residual COH-3/4 cohesin already loaded onto chromosomes (Fig. S6B) may be enough to maintain sister resolution. Unfortunately, we could not test this hypothesis since a longer depletion led to gross morphological aberrations that prevented quantification of sister resolution (Fig. S6A). Alternatively, in the absence of COH-3/4, REC-8 cohesin could be loaded onto chromosomes and perform loop extrusion. Finally, the different effects of NIPBL^SCC-2^ *versus* COH-3/4 depletion on sister resolution could be due to hitherto unappreciated effects of NIPBL^SCC-2^ on REC-8 cohesins, e.g., repositioning them along chromosomes.

Lastly, we analyzed worms lacking Wapl^WAPL-1^, which mediates one of the pathways that removes cohesins from chromosomes (Kueng *et al*. 2006; Gandhi *et al*. 2006). *wapl-1 him-8* worms exhibited a mean DNA:EdU overlap of 52%, not significantly different from *him-8* controls (Fig. S5; unpaired t-test, p=0.91). This indicates that Wapl^WAPL-1^-mediated cohesin unloading is not necessary for sister resolution, consistent with the mild meiotic phenotypes of *wapl-1* null worms (Crawley *et al*. 2016).

## Conclusions

Our work demonstrates that sister chromatids in *C. elegans* meiocytes are resolved from early on in meiotic prophase. Sister resolution occurs both in aligned and unaligned chromosomes (i.e., in the presence of two tethered sisters or four sisters, two from each of the aligned homologs; Fig. 1 and 2), and in both the transcriptionally active autosomes and mostly silenced X chromosome (Fig. 3A-B). We found that sisters are already resolved in early pachytene, only a few hours after they emerge from the replication fork (Fig. 3A-B), suggesting that de-mixing occurs relatively quickly rather than gradually throughout meiotic prophase.

Meiotic sister resolution did not require condensins, nor was it affected by removing the cohesin unloader Wapl^WAPL-1^, REC-8 cohesin or COH-3/4 cohesin (Fig. 3B and S5). However, the cohesin loader NIPBL^SCC-2^ was required for sister resolution (Fig. 4). While the exact NIPBL^SCC-2^-loaded cohesin population relevant for sister resolution remains to be identified, our findings suggest a role for loop extrusion in sister resolution. Given the conservation of meiotic chromosome architecture and of many molecular players, including the cohesin loader NIPBL^SCC-2^, it is likely that our findings in nematodes are relevant across eukaryotes.

Cells regulate homology-directed DSB repair to selectively use the sister or the homolog as a template (Kadyk and Hartwell 1992; Schwacha and Kleckner 1994; Johnson and Jasin 2001; Bzymek *et al*. 2010). Since only inter-homolog exchanges promote chromosome segregation into the gametes during the first meiotic division, it is not surprising that meiotic repair is biased towards the homolog (Lao and Hunter 2010; Zickler and Kleckner 2023). For instance, in worms that experience a single DSB, repair almost always yields a crossover, entailing that the homolog is strongly preferred to the sister as a template for repair (Rosu *et al*. 2011). Nevertheless, sister repair is possible during meiosis (Goldfarb and Lichten 2010; Almanzar *et al*. 2021; Toraason *et al*. 2021), and, based on analysis in budding yeast, homolog bias was surmised to be a result of an active process (Niu *et al*. 2009; Goldfarb and Lichten 2010; Lao and Hunter 2010; Kim *et al*. 2010; Hong *et al*. 2013). In addition, analysis in vegetatively growing cells revealed that spatial proximity is a crucial determinant of template choice (Agmon *et al*. 2013; Lee *et al*. 2016). These observations suggested models for homolog bias where the sisters are intermingled during meiosis, and so the homologous template present on the sister is closer to the DSB and should be actively avoided. For instance, (Hong *et al*. 2013) assumes that the sister is encountered first during repair and is consequently blocked from serving as a template. While the mechanism that distinguishes the homolog from the sister remains unknown, our work shows that the sisters may be as far away from each other as they are from the homolog when meiotic DSBs are induced and repair is taking place. This conclusion suggests that physical proximity *per se* is unlikely to be crucial for implementing homolog bias. Moreover, our identification of a molecular actor important for sister resolution during meiosis - NIPBL^SCC-2^ - opens the door for future investigations of the roles of sister organization in DSB repair dynamics and in the implementation of homolog bias.

## STAR Methods

### Worm strains and growing conditions

Worms were cultured as in (Brenner 1974). Worms used for experimental analysis were age-matched to 24 hours post-L4 and grown at 20ºC. 1mM auxin plates were used for depletion experiments and prepared as in (Zhang *et al*. 2015). All auxin-inducible degron strains used in this study contained the *TIR1* ubiquitin ligase driven by the *sun-1* promotor (Zhang *et al*. 2015). Worms were cultured on NGM plates seeded with OP50 bacteria and moved to auxin plates seeded with OP50 bacteria for the indicated duration of depletion. For analysis of *rec-8*, homozygous progeny of heterozygous mothers were analyzed. See Key Resources Table for a complete list of worm strains.

### Construction of strains

CRISPR was performed as in (Gordon *et al*. 2021) using the co-injection marker *dpy-10*. See Table S1 for all gRNAs and templates. All constructed strains were confirmed via PCR and Sanger sequencing.

### EdU Labeling

EdU labeling was performed essentially as in (Almanzar *et al*. 2021, 2022). Post-labeling chase times were 18 hours for double-label control experiments and 20 hours for single-label experiments. In both cases, the expected sister labeling was confirmed by identifying four differentially labeled populations (from distal to proximal): single-labeled autosomes, single-labeled X chromosomes, double-labeled autosomes, and double-labeled X chromosomes. We only used nuclei with fully synapsed chromosomes to avoid any potential effects of synaptonemal complex disassembly in the transition from pachytene to diplotene. Time prior to EdU labeling was adjusted to ensure all animals were at the same developmental stage when dissected. For confocal images, we used the Alexa 488 Click-it EdU Cell proliferation Kit (Invitrogen) according to the manufacturer’s instructions. For STED images, we used either the Alexa 594 Click-it EdU Cell proliferation Kit (Invitrogen) according to the manufacturer’s instructions, or replaced the fluorophore in the kit with azide-conjugated STAR635 (Abberior) at final concentrations of 3µM. See Key Resources Table for more details.

### Immunofluorescence

Immunofluorescence was conducted as in (Phillips *et al*. 2009). Primary antibody incubations were done overnight at 4°C. STED microscopy samples were mounted in Abberior liquid antifade mounting media. Confocal samples were mounted in NPG-glycerol. Partial maximum intensity projections are shown throughout, except when noted otherwise.

The following antibodies were used: guinea-pig anti-HTP-3 (1:500; (Hurlock *et al*. 2020)), rabbit anti-SYP-5 (1:500; (Hurlock *et al*. 2020)), rabbit anti-RAD-51 (1:10,000; (Harper *et al*. 2011)), rabbit anti-SYP-2 (1:500; (Hurlock *et al*. 2020)), mouse anti-GFP (1:2,000; Roche), rat anti-OLLAS (1:100; Thermofisher)), Cy3 AffiniPure donkey anti-guinea-pig (1:500; Jackson ImmunoResearch), 488 AffiniPure donkey anti-mouse (1:500; Jackson ImmunoResearch), 488 AffiniPure donkey anti-rabbit (1:500; Jackson ImmunoResearch), 594 AffiniPure donkey anti-guinea-pig (1:200; Jackson ImmunoResearch), 594 AffiniPure donkey anti-mouse (1:200; Jackson ImmunoResearch), goat anti-rabbit STAR RED (1:200; Abberior), goat anti-mouse STAR RED (1:200; Abberior), and donkey anti-guinea-pig STAR RED (1:200; Abberior). DNA dyes included DAPI for confocal, and LIVE 460L conjugate DNA (Abberior) and LIVE 560 conjugate DNA (Abberior) for STED. See Key Resources Table for complete details.

### Image acquisition

Confocal microscopy images were taken on a Zeiss LSM880 with Airyscan processing, using the 63x 1.4 NA oil immersion objective. Images were processed using Zen Blue 3.6 (Zeiss) and Imaris (version 9.9, Oxford Instruments). Confocal visible and invisible channels were aligned during image processing using the Channel Alignment function in Zen Blue with the following conditions: third dimension: Z, Quality: Highest, Registration Method: Translation, Interpolation: Cubic.

2D STED images were acquired as in (Almanzar *et al*. 2023), using Abberior STEDYCON mounted on a Nikon Eclipse Ti microscope body with a 100x 1.45 NA oil immersion objective. Due to poor resolution in the z-axis and photobleaching, acquisition and analysis were limited to a single z-section.

3D STED images were acquired on the Abberior Facility Line mounted on an Olympus IX83 microscope body running the Abberior Imspector acquisition and analysis software. Resolution was adjusted to (X) 40nm x (Y) 40nm x (Z) 65nm. 3D STED was carried out at the core microscopy facility at the Scripps Research Facility in La Jolla, CA.

### Spearman’s rank correlation coefficient (SRCC) calculation

SRCC values were calculated using the EzColocalization module for Image J (version 1.52; (Stauffer *et al*. 2018)). For confocal data, individual X chromosomes were isolated out of the 3D confocal images. The middle slice of the chromosome was determined by manual inspection, and two slices, one above and one below, were exported and used for analysis. Each data point represents the SRCC for a single slice. n=8 nuclei from 4 gonads.

For 3D & 2D STED data, we selected regions of interest (i.e., aligned chromosomes) that matched the following criteria: 1) axis tracks could be identified (indicating that the homologs were on each side of the axis in the xy plane, or, in *him-8* animals, that the chromosomes were unaligned); 2) the region of interest was not overlapping with another chromosome; and 3) the chromosome contained EdU labeling. N2 single-label: 17 chromosomes were selected from 6 different nuclei from 4 different gonads; N2 double-label: 15 chromosomes were selected from 7 different nuclei from 4 different gonads; SCC-2-AID + auxin: 22 chromosomes were selected from 8 different nuclei from 4 different gonads; *him-8*: 10 chromosomes were selected from 3 different gonads.

### Single chromosome masking for 3D STED images

Chromatin masks of single paired chromosomes (as in Figs. 1 and 4) were generated using Imaris (version 9.9, Oxford Instruments). The Surfaces function was used to construct a DNA surface with a smoothing detail of 0.05μm with Absolute Intensity thresholding. The Split Touching Objects function was used with a Seed Diameter of 0.04μm. Individual chromosomes were manually verified.

### Deriving the unlabeled sister in 3D STED images

To computationally derive the unlabeled sister, we used custom code in MATLAB to normalize the DNA and EdU channels to the 90% percentile intensity. We then subtracted the EdU values from the DNA values and normalized the intensity. All channels were smoothed using the imgaussfilt3 function with sigma = 1. MATLAB code available upon request.

### Volumetric quantification of sister resolution in confocal images

To analyze the overlap between the sisters, we used Imaris (version 9.9, Oxford Instruments). Individual nuclei were cropped in 3D from full gonad images where we identified the meiotic stage and confirmed expected EdU labeling. From each nucleus, chromosomes used for quantification were further cropped in 3D. Chromosomes chosen for analysis were: 1) unaligned, identified by the lack of synaptonemal complex central region (SYP-2 or SYP-5 signal) colocalizing with the axis (HTP-3 signal); 2) not overlapping with other chromosomes. Using the EdU and DAPI fluorescence channels, we constructed two surfaces using the Surfaces function in Imaris with default thresholding. Two volumetric ratios were computed using the Export Individual Values Surfaces Ratio function: 1) the ratio of DAPI:EdU and 2) the ratio of EdU:DAPI. Each data point represents a single chromosome.

### Axis length measurements

The pachytene nuclei used for quantification all harbored only fully synapsed chromosomes verified by the overlap of the synaptonemal complex central region marker SYP-5 and the axis marker HTP-3. Imaris (version 9.9, Oxford Instruments) was used to trace HTP-3 (axis) staining and calculate chromosome length using the Measurement Points function.

### Transition zone measurements

Transition zone lengths were calculated by stitching together multiple (typically 2-3) images to generate whole gonad images with DAPI, nucleolus (DAO-5) and axis (HTP-3) staining. The transition zone was denoted by crescent-shaped chromatin morphology and the off-center localization of the nucleolus. We defined the end of the transition zone as the last row with two crescent-shaped nuclei followed by two rows with zero or one crescent-shaped nuclei. Transition zone length was computed by using FIJI (version 2.14) and calculating the overall length of the gonad as the skeletonized distance between the mitotic tip to the end of pachytene (where nuclei become single-file).

### Fluorescence quantification for SCC-2 and COH-3 depletion

Pachytene regions were imaged with consistent settings among samples and slides, including laser power. Z-stacks were acquired and average intensity was projected in Zen Black. Exported TIFF files were opened in ImageJ (version 2.14) and fluorescence intensity of the channel of interest was calculated for nuclei of equal sizes. Fluorescence intensity was normalized to the background signal of an area the same size outside the nuclei.

## Supporting information

Key Resources Table

## Acknowledgements

We thank the Rog lab for discussions and support; Kathy Spencer and Scott Henderson at the Scripps Research Facility for help with the 3D-STED; Enrique Martinez-Perez for sharing the *scc-2-aid* strain prior to publication; Weston Stauffer for advice on the open-source ImageJ EzColocalization plugin module; Sara Nakielny for comments on the manuscript and editorial work; and Yumi Kim for antibodies. Some strains used in this work were provided by the CGC, which is funded by NIH Office of Research Infrastructure Programs (P40 OD010440). We acknowledge the HSC Cell Imaging Core at the University of Utah for use of the STEDYCON microscope. AH is supported by Developmental Biology Training Grant T32HD007491 from NICHD. Work in the Rog lab is supported by grant R35GM128804 from NIGMS.

