## Supplemental Figures and Legends for "Early resolution of sister chromatids during *C. elegans* meiosis"

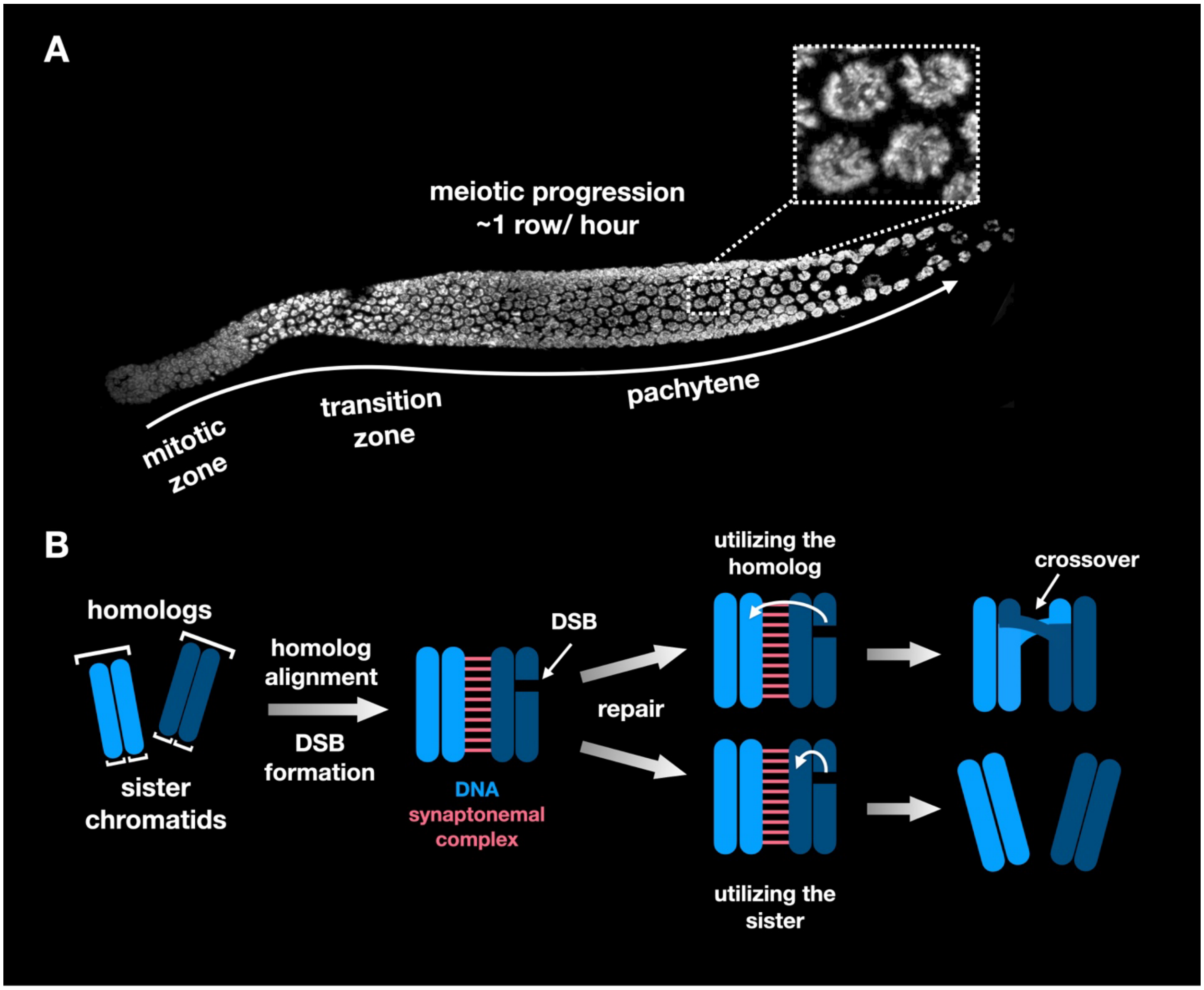


**Fig. S1**

**Figure S1: Gonad organization in *C. elegans*, related to Figure 1**

A. Whole gonad stained with DAPI (white), with meiosis progressing from left to right. Nuclei undergo mitotic division and then enter meiosis in the so-called 'transition zone', characterized by crescent-shaped nuclear morphology (equivalent to the classically defined leptotene and zygotene stages). Nuclei move at a rate of ~1 row per hour through the gonad into pachytene, diplotene, and eventually diakinesis, where chromosomes become hyper-condensed and are about to undergo the first meiotic division (not shown in this gonad). Nuclei in pachytene are shown in the zoomed-in view.

B. During meiotic progression, each nucleus contains six pairs of homologous chromosomes (homologs), one set from each parent (for clarity, only a single pair is shown). These similar homologs each contain a genetically-identical pair of sister chromatids. The chromosomes pair and align with their homolog and assemble the synaptonemal complex (pink) at the interface between them. A meiosis-specific endonuclease (Spo11; SPO-11 in worms; (Keeney *et al.* 1997; Dernburg *et al.* 1998)) creates DSBs that can be repaired in a variety of ways. Two possible flavors of homologous recombination - the prominent meiotic repair pathway - are shown. The first (top), utilizes the other homolog as a template, allowing for formation of crossovers that enable correct chromosome segregation in the first meiotic division. Utilizing the sister chromatid as a template (bottom) does not generate crossovers. Chromosomes lacking a crossover segregate randomly, leading to aneuploid gametes and progeny.

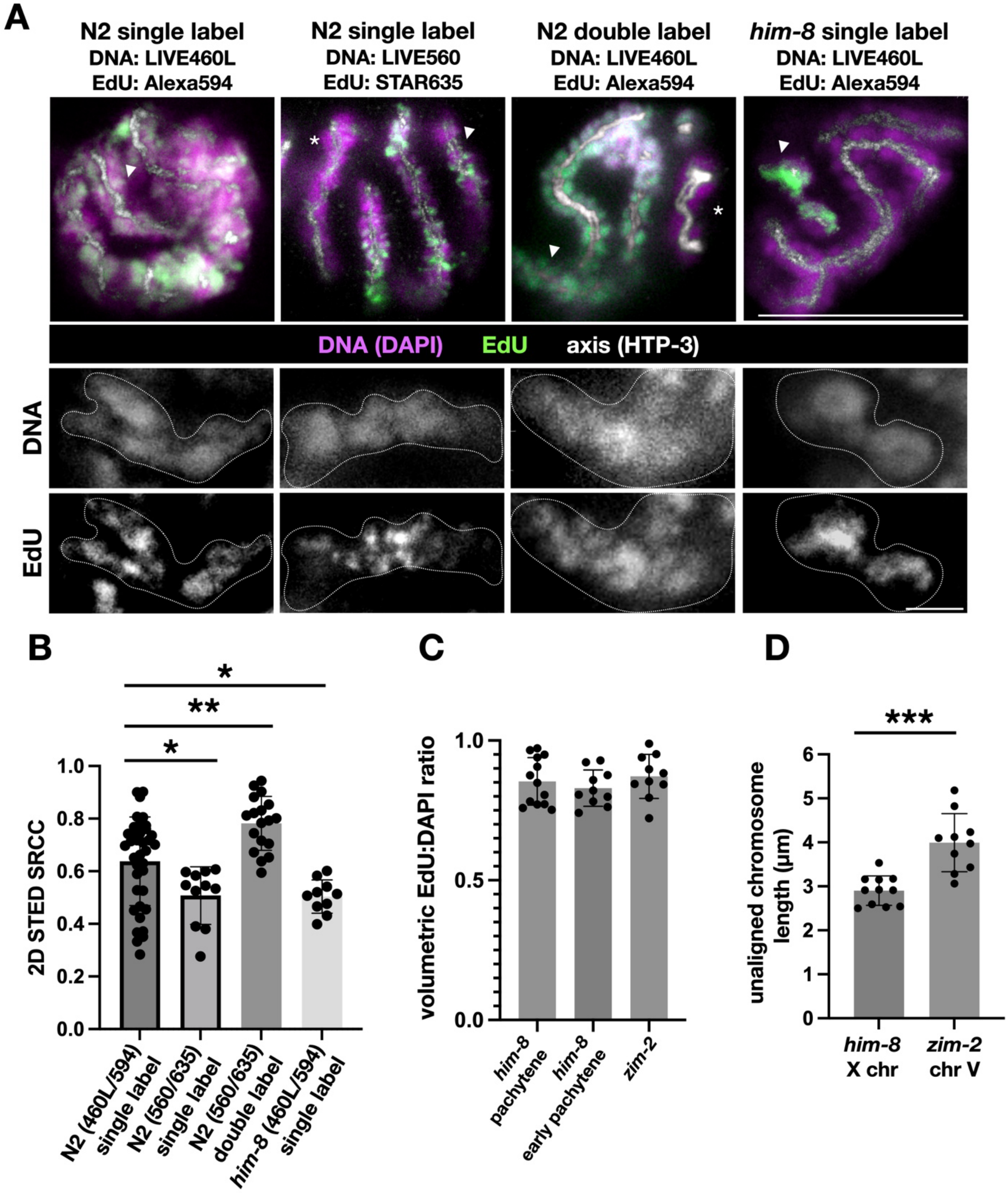
A
N2 single label
DNA: LIVE460L
EdU: Alexa594
N2 single label
DNA: LIVE560
EdU: STAR635
N2 double label
DNA: LIVE460L
EdU: Alexa594
him-8 single label
DNA: LIVE460L
EdU: Alexa594
DNA (DAPI)
EdU
axis (HTP-3)
DNA
EdU
B
2D STED SRCC
N2 (460L/594) single label
N2 (560/635) single label
N2 (560/635) double label
him-8 (460L/594) single label
C
volumetric EdU:DAPI ratio
him-8 pachytene
him-8 early pachytene
zim-2
D
unaligned chromosome length (μm)
him-8 X chr
zim-2 chr V
***

Fig. S2

**Figure S2: Additional analysis relating to Figures 1, 2 and 3**

A. Top, 2D-STED images (single z-section) of wild-type (N2) animals with single- and double-labeled chromosomes (DNA:LIVE460L, EdU: Alexa594), wild-type (N2) single label animals with an alternative dye combination (DNA: LIVE560, EdU: STAR635), and *him-8* animals with the unpaired X chromosome (white arrowhead; DNA:LIVE460L, EdU: Alexa594). Note that the 3D-STED images in Fig. 1 used LIVE460L to label DNA and Alexa594 to label EdU. Scale bar = 5μm. Asterisks represent unlabeled X chromosomes. Second and third row, single-channel greyscale images of the regions of interest from the top row (indicated by arrowhead), with the outline of the chromosome shown with a dashed white line.

B. Spearman's rank correlation coefficient (SRCC) between the DNA and EdU channels from 2D-STED single-labeled N2 worms (DNA: LIVE460L; EdU: Alexa594), single-labeled N2 worms (DNA: LIVE560; EdU: STAR635), double-labeled N2 worms (DNA: LIVE460L; EdU: Alexa594), and unpaired X chromosomes from single-labeled *him-8* worms (DNA: LIVE460L; EdU: Alexa594; n = 37,11,18 and 11, respectively). One-way ANOVA with multiple comparisons: N2 single (594) vs. N2 single (635) p=0.02; N2 single (594) vs. N2 double (594) p=0.0014; N2 single (594) vs. *him-8* single (594) p=0.023.

C. Volumetric EdU:DAPI ratio for single-labeled *him-8* pachytene, *him-8* early pachytene and *zim-2* worms (n=13, 10, and 10, respectively), related to the ratios in Fig. 3B. One-way ANOVA with multiple comparisons: *him-8* pachytene vs. *him-8* early pachytene p=0.96; *him-8* pachytene vs. *zim-2* p=0.99.

D. Length of the chromosome axis of the unaligned X chromosome in *him-8* animals and of the unaligned chromosome V in *zim-2* animals (n = 11 and 12 respectively). Unpaired t-test p=0.0007. The X chromosome is 18.1Mbp and chromosome V is 20.8Mbp. Rather than the expected 15% increase in chromosome length, chromosome V was 30% longer than the X chromosome.

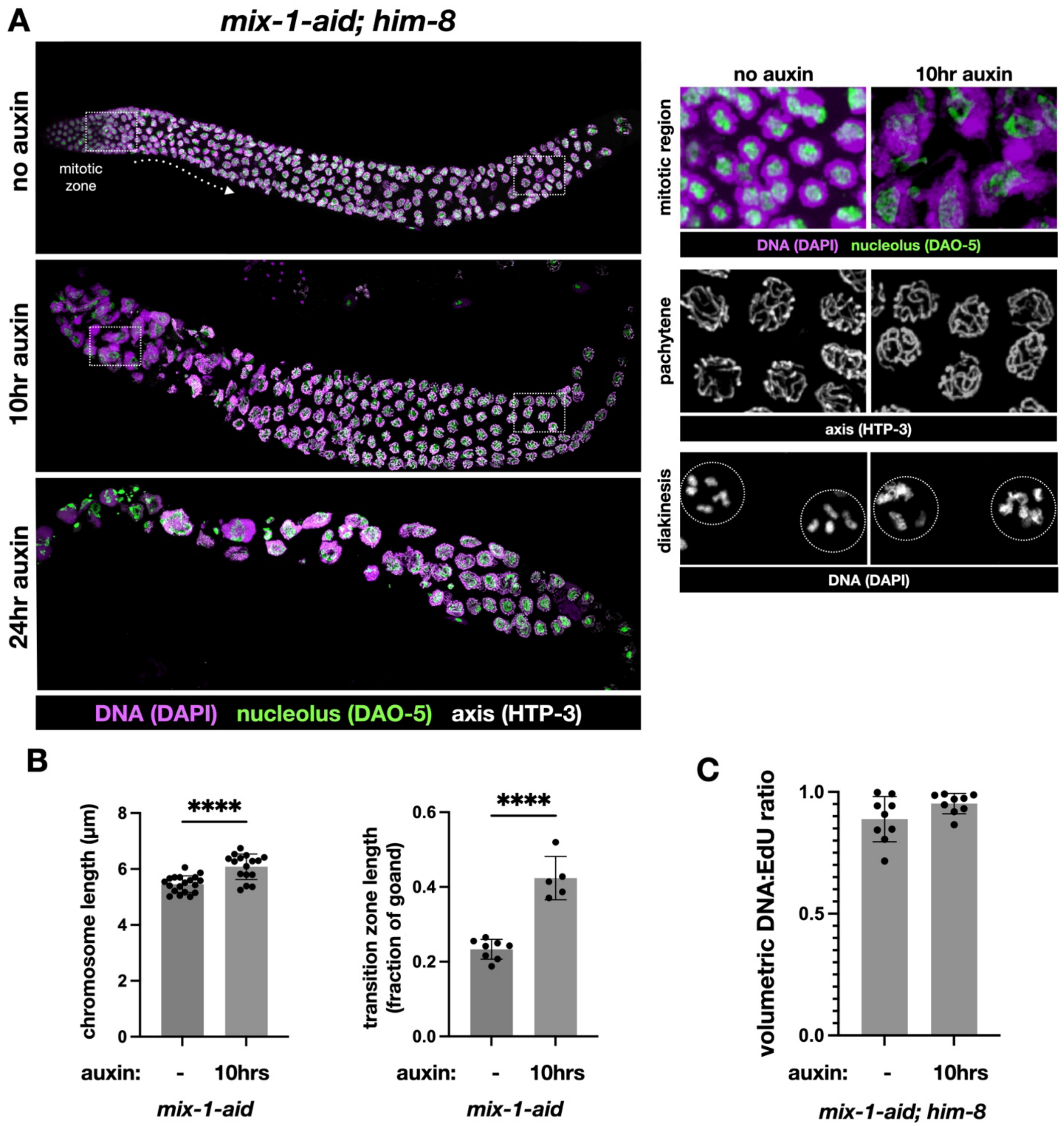
A
mix-1-aid; him-8
no auxin
mitotic zone
10hr auxin
24hr auxin
DNA (DAPI) nucleolus (DAO-5) axis (HTP-3)
no auxin
10hr auxin
mitotic region
DNA (DAPI) nucleolus (DAO-5)
pachytene
axis (HTP-3)
diakinesis
DNA (DAPI)
B
chromosome length (µm)
****
auxin: - 10hrs
mix-1-aid
transition zone length (fraction of goand)
****
auxin: - 10hrs
mix-1-aid
C
volumetric DNA:EdU ratio
auxin: - 10hrs
mix-1-aid; him-8

Fig. S3

**Figure S3: Additional characterization of condensin depletion, related to Figure 3**

A. Left, whole gonad from *mix-1-aid; him-8* animals without auxin (control; top), after 10-hour depletion (middle), and after 24-hour depletion (bottom; confocal image projections). DNA (DAPI), magenta; nucleolus (DAO-5), green; axis (HTP-3), green. Zoomed-in views on the right highlighting the mitotic region (top), pachytene (middle), and DAPI staining bodies (bottom) in control (left) and 10-hour depletion (right). Note the large and disorganized mitotic nuclei upon auxin treatment, indicative of mitotic failure, as well as the incompletely condensed chromosomes prior to the first meiotic division (diakinesis). Importantly, the morphology of pachytene nuclei is not visibly affected.

B. Length of the chromosome axis and length of the transition zone in control (n=19 & n=8) and 10-hour depleted worms (n=16 & n=5). Unpaired t-test $p<0.0001$ for both metrics. The elongated chromosome axis and extended transition zone are consistent with the phenotypes exhibited by condensin hypomorphs (Mets and Meyer 2009).

C. Volumetric EdU:DAPI ratio for single label *mix-1-aid* control and 10-hour depleted (n=9 for both samples) worms, related to Fig. 3D. Unpaired t-test p=0.079.

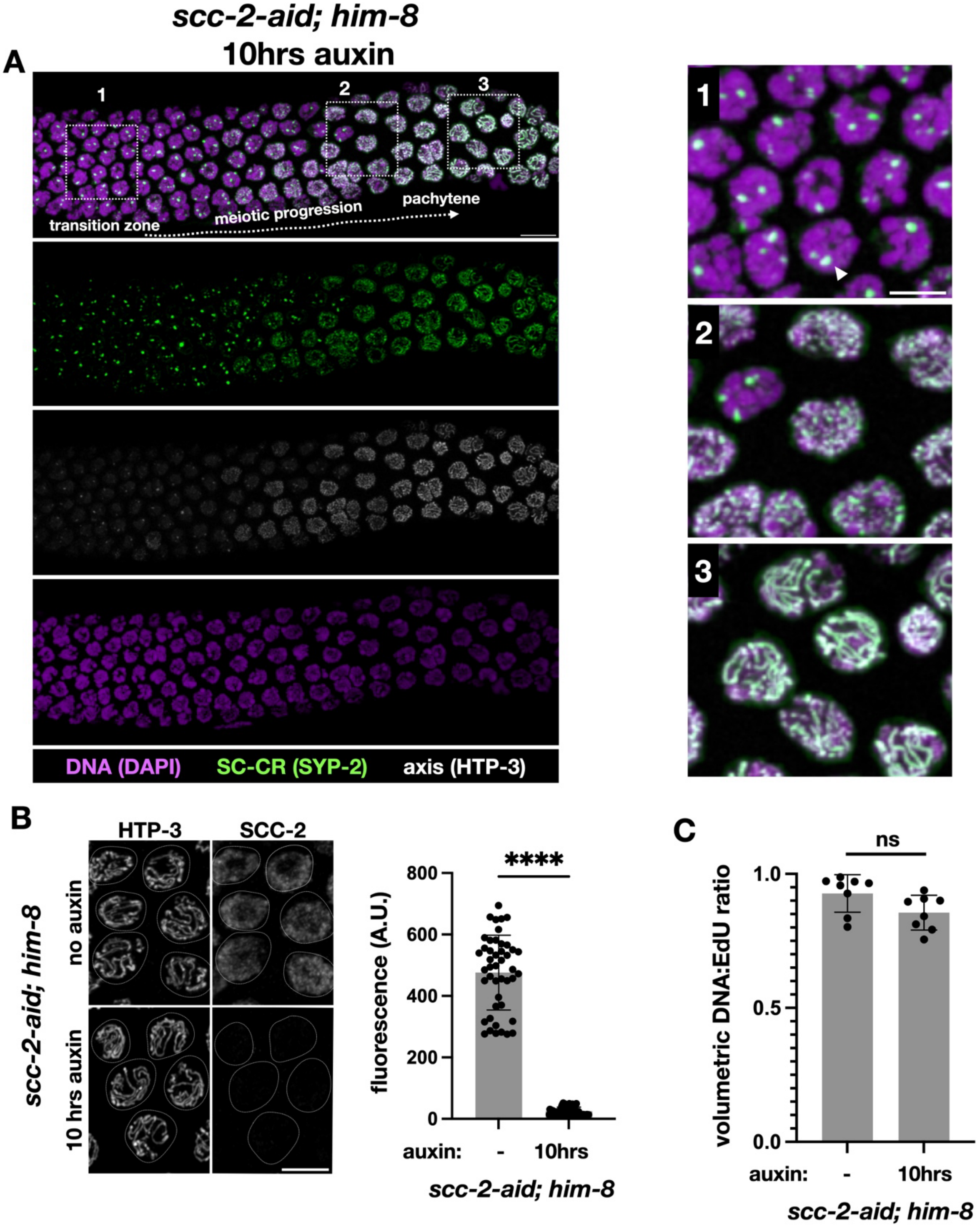
scc-2-aid; him-8
10hrs auxin
A
1
2
3
transition zone
meiotic progression
pachytene
DNA (DAPI)
SC-CR (SYP-2)
axis (HTP-3)
B
HTP-3
SCC-2
scc-2-aid; him-8
no auxin
10 hrs auxin
****
800
600
400
200
0
fluorescence (A.U.)
auxin:
-
10hrs
scc-2-aid; him-8
C
ns
1.0
0.5
0.0
volumetric DNA:EdU ratio
auxin:
-
10hrs
scc-2-aid; him-8

Fig. S4

**Figure S4: Additional characterization of NIPBL$^{SCC-2}$ depletion, related to Figure 4**

A. Left, pachytene region from gonads of *scc-2-aid; him-8* worms after 10 hours on auxin (confocal image projections). Top, merged image with DNA (DAPI) in purple, synaptonemal complex central region (SC-CR; SYP-2) in green, and axis (HTP-3) in white. Individual channels are shown below. Scale bar = 10 µm. Right, zoomed-in views. (1) In transition zone nuclei, the synaptonemal complex central region forms aggregates, called polycomplexes (arrowhead), as has been shown in *scc-2* null worms (Lightfoot *et al.* 2011). (2) Nuclei in the pachytene region where the axis and the synaptonemal complex central region show a punctate pattern, indicative of disassembly. (3) Pachytene nuclei just a few rows later in the gonad, where the axis and synaptonemal complex central region are still intact. Nuclei in region 3 are representative of nuclei used for quantification in Fig. 4. Scale bar = 5µm.

B. Pachytene nuclei from *scc-2-aid; him-8* worms without auxin and after 10 hours on auxin labeled with an axis marker (HTP-3) as a reference and NIPBL$^{SCC-2}$ (SCC-2; using antibodies against GFP). Scale bar = 5µm. Right, fluorescence in no auxin and in 10-hour depleted worms (n=45 and 72, respectively). Unpaired t-test $p<0.0001$.

C. Volumetric EdU:DAPI ratio for single-labeled *scc-2-aid* control and 10-hour depleted worms (n=8 for each sample). Unpaired t-test $p=0.051$. Related to Fig. 4B.

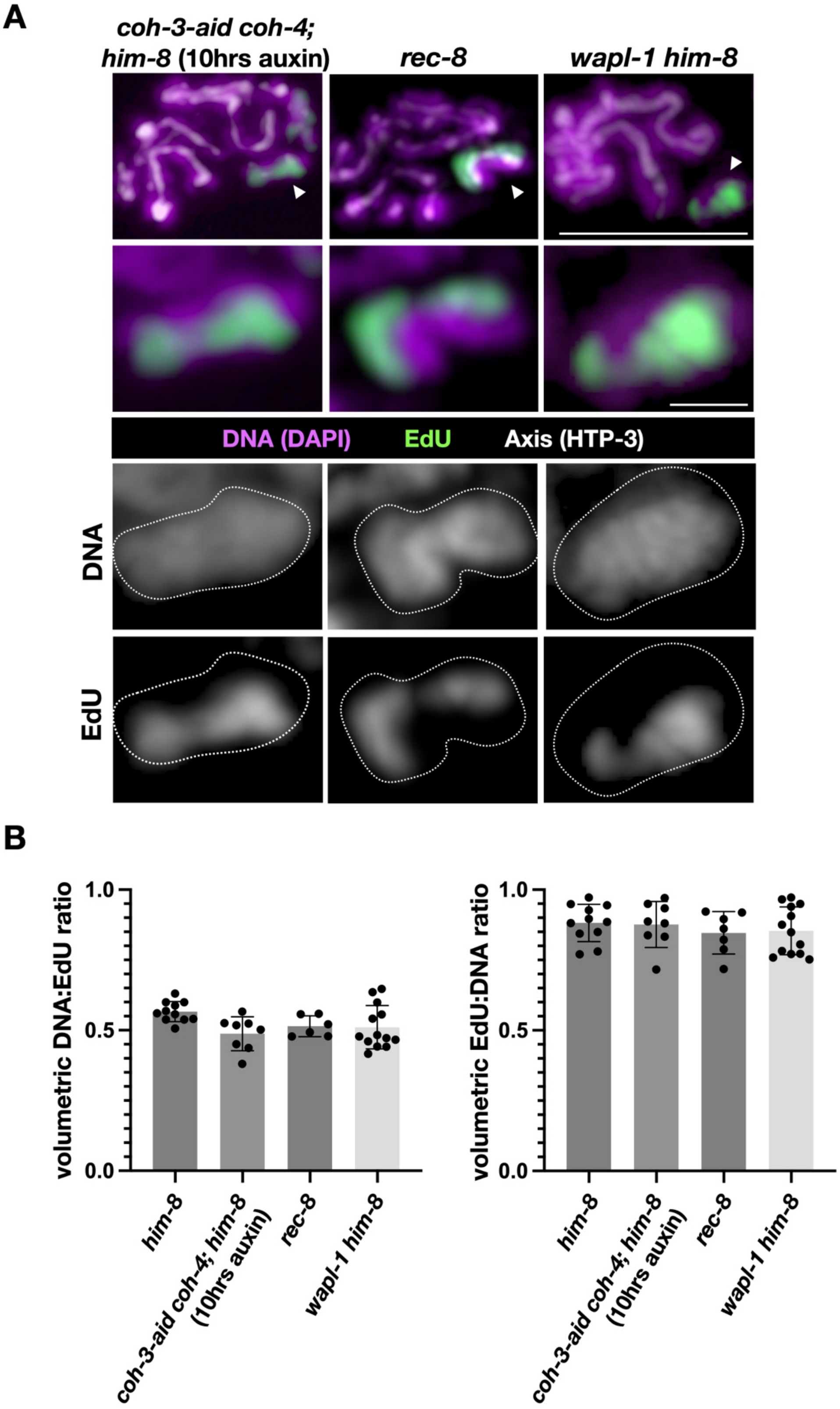
A
coh-3-aid coh-4; him-8 (10hrs auxin)
rec-8
wapl-1 him-8
DNA (DAPI)
EdU
Axis (HTP-3)
DNA
EdU
B
volumetric DNA:EdU ratio
1.0
0.5
0.0
him-8
coh-3-aid coh-4; him-8 (10hrs auxin)
rec-8
wapl-1 him-8
volumetric EdU:DNA ratio
1.0
0.5
0.0
him-8
coh-3-aid coh-4; him-8 (10hrs auxin)
rec-8
wapl-1 him-8

**Fig. S5**

**Figure S5: Additional characterization of the effects of cohesins on sister resolution, related to Figure 4**

A. Top, confocal images of pachytene nuclei from 10-hour auxin depleted *coh-3-aid coh-4 him-8, rec-8,* and *wapl-1 him-8* worms. DNA (DAPI), purple; EdU, green; axis (HTP-3), white. Second row, zoomed-in view of the unaligned, EdU-labeled X chromosome. Based on EdU incorporation, the EdU-labeled chromosome shown in rec-8 worms is also the X chromosome. Third and fourth rows, single-channel greyscale images, with the outline of the chromosome shown (dashed white line). Scale bar is 5µm for the whole nucleus and 1µm for the single chromosome.

B. Volumetric DNA:EdU and EdU:DNA ratios of *him-8* (from Fig. 2), 10-hour auxin *coh-3-aid coh-4 him-8, rec-8* and *wapl-1 him-8* worms (n=13, 11, 8 and 6, respectively). ANOVA p values are all not significant.

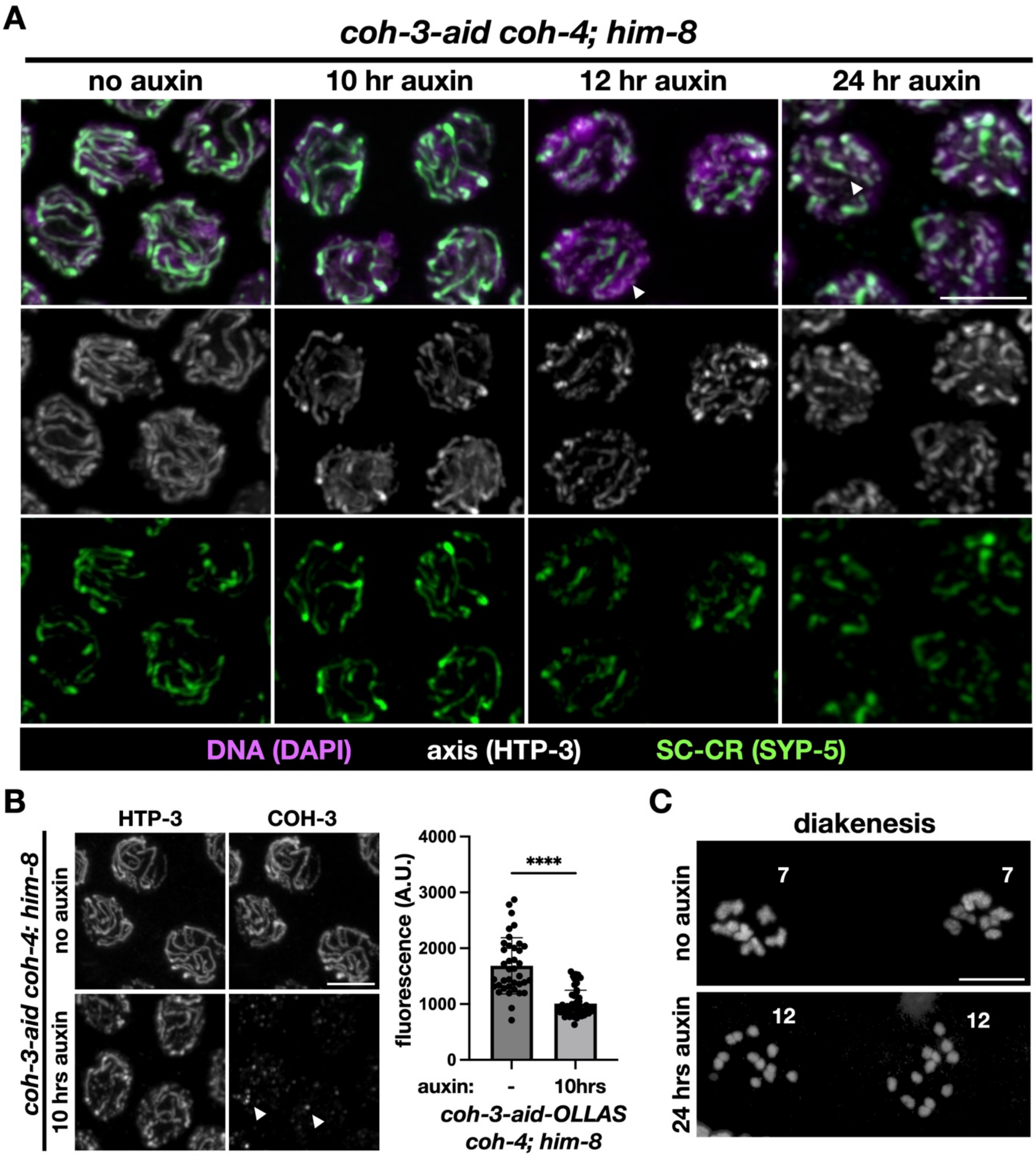
A
coh-3-aid coh-4; him-8
no auxin
10 hr auxin
12 hr auxin
24 hr auxin
DNA (DAPI)
axis (HTP-3)
SC-CR (SYP-5)
B
coh-3-aid coh-4; him-8
HTP-3
COH-3
no auxin
10 hrs auxin
4000
3000
2000
1000
0
fluorescence (A.U.)
****
auxin:
-
10hrs
coh-3-aid-OLLAS
coh-4; him-8
C
diakenesis
no auxin
24 hrs auxin
7
7
12
12

**Fig. S6**

**Figure S6: Characterization of COH-3/4 depletion, related to Figure 4**

A. Confocal images of pachytene nuclei from *coh-3-aid coh-4; him-8* worms. DNA (DAPI), purple; synaptonemal complex central region (SC-CR; SYP-5), green; axis (HTP-3), white. Scale bar = 5µm. Worms were either not treated with auxin or treated with auxin for 10, 12 or 24 hours. In no auxin control worms, the axis and synaptonemal complex central region overlap, indicating homologs are paired and aligned (except the X chromosome, which is unaligned due to the absence of *him-8*). After 10-hour depletion, the axis and synaptonemal complex central region are still mostly intact, allowing us to quantify sister resolution. However, after 12-hour depletion, the axis staining becomes punctate, indicating the axes started to fall apart, and the synaptonemal complex central region staining becomes not continuous. These effects on chromosome organization prevented reliable quantification of sister resolution. These phenotypes became worse after 24 hours of depletion.

B. Pachytene nuclei from *coh-3-aid-OLLAS coh-4; him-8* worms either not treated with auxin or after 10-hour depletion. The axis (HTP-3) serves as a reference channel; COH-3 is labeled using antibodies against the OLLAS tag. Note the residual COH-3 after 10-hour depletion (white arrowheads). Scale bar = 5µm. Right, COH-3 (OLLAS) fluorescence in no auxin and 10-hour depleted worms (n=38 and 28, respectively). Unpaired t-test $p<0.0001$.

C. Diakinesis nuclei from no auxin and 24-hour depleted *coh-3-aid coh-4; him-8* worms. Each nucleus in no auxin worms has 7 DAPI bodies, as expected (5 paired autosomes and 2 unpaired X chromosomes), compared to the 12 DAPI staining bodies after 24 hours on auxin, consistent with the *coh-3 coh-4* null phenotype (Severson and Meyer 2014).
