## Supplementary material for "Early resolution of sister chromatids during *C. elegans* meiosis": Key Resources Table

| REAGENT or RESOURCE | SOURCE | IDENTIFIER |
|---|---|---|
| Antibodies | | |
| Guinea pig anti-HTP-3 | Yumi Kim Lab | n/a |
| Rabbit anti-SYP-2 | Yumi Kim Lab | n/a |
| Rabbit anti-SYP-5 | Yumi Kim Lab | n/a |
| Mouse anti-DAO-5 | DSHP | ID: AB_10573805 |
| Mouse anti-GFP | Roche | #11814460001 |
| Rat anti-OLLAS | invitrogen | MA5-16125 |
| Cy3 AffiniPure Donkey anti-guinea pig IgG | Jackson Immuoresearch | Cat#706-165-148; RRID: AB_2340460 |
| 647 AffiniPure Donkey anti-rabbit IgG | Jackson Immuoresearch | Cat#711-605-152 RRID: AB_2492288 |
| Cy3 AffiniPure Donkey anti-rat IgG | Jackson Immuoresearch | Cat#712-165-153 RRID: AB_2340667 |
| 488 AffiniPure Donkey anti-mouse IgG | Jackson Immuoresearch | Cat#715-545-150 RRID: AB_2340846 |
| Abberior STAR RED Goat anti-guinea pig IgG | Abberior | Item #STRED |
| Abberior LIVE 460L-DNA conjugate | Abberior | Item #LV460L |
| Abberior STAR 635-azide conjugate | Abberior | Item #ST635 |
| Abberior LIVE 560 DNA | Abberior | Item #LV560 |
| Abberior STAR 460L goat anti-guinea pig | Abberior | Item #ST460L |
| Chemicals, peptides, and recombinant proteins | | |
| Indole-3-acetic acid (auxin) | VWR | Cat#AAA10556-36 |
| Cas9 nuclease V3 | IDT | Cat#1081058 |
| Abberior MOUNT | Abberior | Item #MM |
| Critical commercial assays | | |
| Click-it EdU Cell proliferation Kit, Alexa Fluor 488 dye | Invitrogen | Cat #C10337 |
| Click-it EdU Cell proliferation Kit, Alexa Fluor 594 dye | Invitrogen | Cat #10339 |
| Experimental models: Organisms/strains | | |
| *C. elegans* N2 | CGC | N2 |
| *him-8(tm611) IV* | CGC | CA257 |
| *mix-1(slc15[mix-1::aid]) II; ieSi38[sun1p::tir1::mRuby::sun-1 3'UTR+ Cbr-unc-119(+)] him-8(tm611) IV* | This study | ROG386 |
| *mix-1(slc15[mix-1::aid]) II; ieSi38[sun1p::tir1::mRuby::sun-1 3'UTR+ Cbr-unc-119(+)] IV* | This study | ROG391 |
| *scc-2(fq23[scc-2::aid::gfp])II; ieSi38[sun1p::tir1::mRuby::sun-1 3'UTR+Cbr-unc-119(+)] IV* | Enrique Martinez-Perez Lab | ATG282 |
| *scc-2(fq23[scc-2::aid::gfp])II; ieSi38[sun1p::tir1::mRuby::sun-1 3'UTR+Cbr-unc-119(+)] him-8(tm611) IV* | This study | ROG447 |
| *zim-2(tm574) IV* | CGC | CA258 |
| *rec-8(ok978) IV/nT1 [qIs51] (IV;V)* | CGC | VC666 |

| | | |
|---|---|---|
| *coh-3(slc16[coh-3::aid::OLLAS]) coh-4[slc17] V; ieSi38[sun1p::tir1::mRuby::sun-1 3'UTR+ Cbr-unc-119(+)] him-8(tm611) IV* | This study | ROG446 |
| *wapl-1(slc18) him-8(tm1611) IV* | This study | ROG481 |
| Software and algorithms | | |
| Zen Blue | Carl Zeiss Microscopy | version 3.9 |
| Zen Black | Carl Zeiss Microscopy | version 2.3 |
| Prism | GraphPad | version 10.0 |
| Imaris | Oxford Instruments | version 9.9 |
| ImageJ | | version 1.52 |
| FIJI | | version 2.14 |
| Abberior Light Box IMSPECTOR | Abberior | 2024 free version |
| MATLAB | MathWorks | R2024a |
| Other | | |
| LSM 880 confocal microscope with AiryScan | Carl Zeiss Microscopy | |
| Abberior STEDYCON | Abberior | |
| Abberior Facility line STED | Abberior | |
